# Patterning High to Low Heterogenous Fluid Shear Stress Landscapes with Multiphoton Inner Laser Lithography (MILL) for Live Cell Adhesion and Translocation

**DOI:** 10.1101/2022.06.17.496569

**Authors:** Yean J. Lim, Junxiang Zhang, Hanqi Lin, Tienan Xu, Yongxiao Li, Zhiduo Zhang, Sarah M. Hicks, Ivan Chudinov, Dmitry Nechipurenko, Elizabeth E. Gardiner, Woei M. Lee

## Abstract

Heterogenous fluid shear stress is known to provide mechanical cues for cell adhesion and translocation. To assemble 3D microstructures using current fabrication methods in a single channel and recapitulate in vivo heterogenous fluid flow would require hours of fabrication and specialized equipment. Inspired by the traditional art form of inside painting, we developed a technique for 3D fabrication of micro-patterned flow channels and mixed in vivo fluid flow in a matter of minutes. We termed this technique Multiphoton Inner Laser Lithography (MILL). We further showed that when combined with adaptive optics, MILL is compatible with both flat and curved channel shapes. MILL recapitulated in vivo tissue topology and 3D fluid flow within tissue stroma (low fluid shear, 0 – 3.5 dynes/cm^2^) and blood vessel (high fluid shear, 0 – 80 dynes/cm^2^). We demonstrate fibroblast cell and platelets adhere and translocate differently between laminar flow patterns that are homogenous versus heterogeneous in real time. Parallel strips of MILL channels were assembled for simultaneous platelet function test to quantify the efficacy of an antithrombotic GPVI Fab (~2000 microthrombi per test). The MILL technique can be readily reproduced *in vivo* fluid flow in minutes and benefit preclinical screening of drug pharmacokinetics.

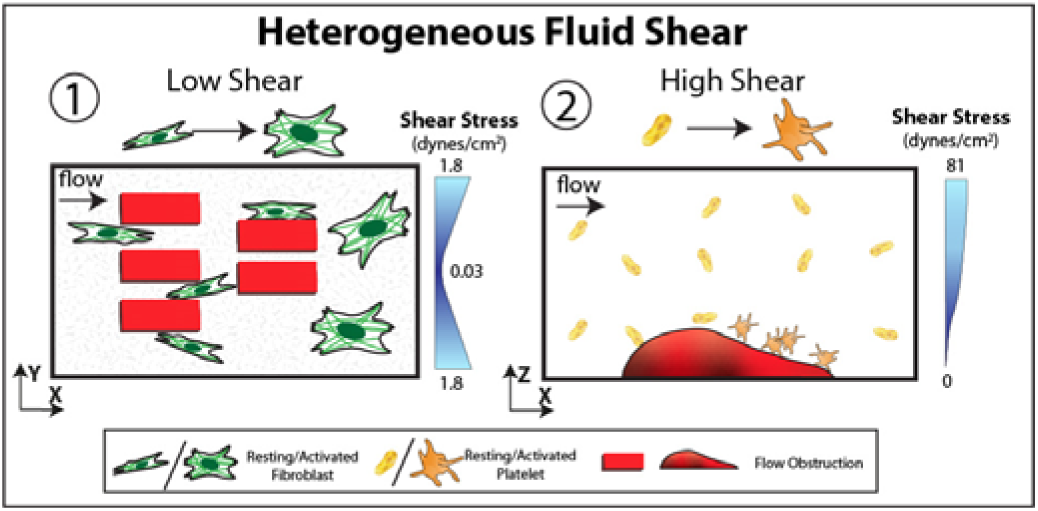

**Significant points:**

- *MILL channels* are made from commercially available off the shelf components using a standard multiphoton imaging system
- Varying degrees of *in vivo* heterogenous laminar flow is shown to directly influence cells translocating on thinly coated collagen surfaces
- Parallel strips of *MILL* channels were assembled for simultaneous platelet function tests (~2000 microthrombi per test).

## Introduction

Extracellular fluid flowing through porous tissue matrix and blood vessels generates an uneven distribution of laminar flow rates and imposes heterogenous fluid shear stress on surrounding adherent cell population. Laminar flows are often assigned a single shear rate ^1^ because it exhibits a parabolic profile across a uniform smooth channel. *In vivo* flow through living tissue encounters multiple barriers (e.g. surface protrusions, cells and extracellular matrix) that alter the parabolic flow, creating different degrees of laminar flow in the interstitium and blood vessels. Hence, ‘laminar flow’ does not fully encompass *in vivo* flow complexity ^2^. Fluid flowing from the heart to interstitial tissue generates a range of laminar fluid shear landscapes ranging from 0 – 80 dynes/cm^2^. We propose that laminar flow landscapes possess mixed laminar shear which can be referred to as ‘heterogeneous flow’. Simple flow landscapes with fixed laminar shear are therefore referred as ‘homogeneous flow’. While it has been shown that *in vivo* heterogenous fluid flow varies from the low fluid shear regimes (0 – 3.5 dynes/cm^2^) in interstitial tissue matrix to high fluid shear regimes (0 – 80 dynes/cm^2^) in vasculature ^3^, there has been limited studies into *in vivo* fluid motion that regulate cell adhesion and migration ^2,4,5^. We argue that this is also because of lack of direct 3D flow channel fabrication methods to adequately reflect complex heterogeneous *in vivo* flow environment.

All cells have evolved mechanisms to sense and navigate through heterogenous fluid shear landscapes using mechanosensitive signaling receptors such as Piezo and transient receptor potential (TRP) family proteins ^3,6^. More prominently, endothelial cells ^3^, fibroblast ^6^, platelets ^7^ and immune cells ^8^ residing in these landscapes require Piezo or TRP to migrate and activate. Heterogeneous fluid shear landscapes are responsible for triggering intracellular signaling, transduction^9^ and cell migration^10^ and are important in maintaining the homeostasis of tissue niches ^11^, initiating tissue repair ^12^, bone development ^13^, cancer metastasis ^14,15^, angiogenesis ^4^, regulating cardiac fibrosis and impacts on thrombosis ^16^. Despite this, the majority of tissue-on-chip ^17^ or organ-on-chip ^18^ systems are generally designed with homogeneous fluid shear stress (FSS) in soft lithographic hollow channels lined by a single layer of cultured cells or tissue. Complex 3D microstructures fabricated in a single microfluidic channel are necessary to reproduce a mixture of fluid shear conditions present in *in vivo* microenvironments. Common fabrication methods to create microstructure patterning in a single flow channel either uses soft lithography (Figure 1 A) i)) or fused deposition (Figure 1 A) ii)). Both of the techniques require multiple stepwise deposition of curable resin or polydimethylsiloxane (PDMS) respectively, onto open two-dimensional surfaces ^19^ that often require hours of preparation and specialized instruments.

**Figure 1.**
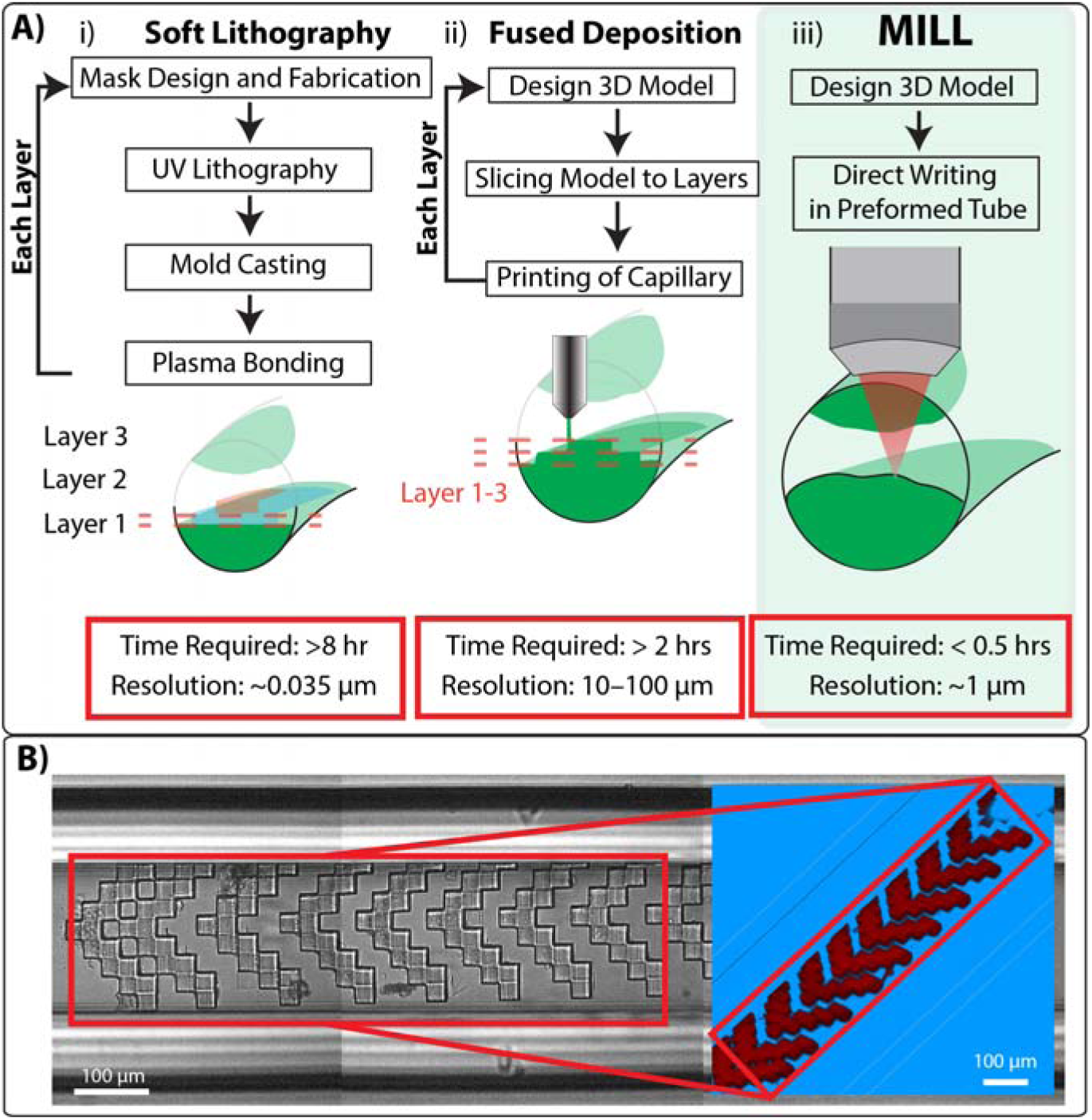
Current approaches for fabricating microfluidic channels with 3D patterning. **A)** Diagram of steps and time requirement to fabricate a circular microchannel with an irregular-shaped structure using (i) soft lithography, (ii) fused deposition modeling or (iii) MILL. The lithography resolutions and time incurred are indicated below each process. **B)** Brightfield image and 2-photon fluorescence image (inset) of herringbone structure in a square glass capillary tube.

Aside from flexibility in 2D micropatterning, high optical transparency (>90%) and surface functionalization are essential in molecular and cellular assays ^20^ for genomics, proteomics and phenotypic quantification ^21^. There are significant fabrication hurdles to develop 2D soft lithography techniques ^22^ for fully enclosed 3D surfaces that has high optical transparency. (1) Soft lithography processes are optimized for open and exposed flat surfaces that require passivating steps ^23^, (2) soft lithography requires specialized equipment (UV lamp, thermal heating oven, desiccator, plasma cleaner) ^24^, (3) PDMS variably absorbs different hydrophobic molecules and so confound studies involving the delivery of small molecules into cells ^25^. Taken together, standard soft lithography involves hours of labor-intensive, repeated set of steps (Figure 1A) ^26^ to achieve 3D microstructures. To advance cell biology questions incorporating complex heterogeneous *in vivo* flow environment, we propose to develop a full 3D lithography method ^27,28^ that is directly compatible with enclosed 3D flow channels ^29^. The goal of direct 3D modification in an enclosed flow channel is to mimic *in vivo* flow conditions of different microenvironments. In the tissue microenvironment, the 3D surface roughness influences the trajectory of fluid flow ^2,5^ that is important in platelet production ^1^, transmural nutrient delivery ^30^ atherosclerosis ^5^ and tumor metastasis ^14^.

In this study, we describe a direct 3D fabrication method, Multiphoton Inner Laser Lithography (MILL), that modifies glass capillaries ^31,32^ for use in a microfluidic flow assay within 1 hour (Figure 1A, iii) with micrometer resolution. In doing so, we, for the first time, gain access to *in vivo*, 3D heterogenous fluid flow landscapes present in vasculature and stroma. MILL is inspired by the art form of inside painting ^33^, that uses multiphoton absorption to sculpt the inner surfaces of transparent glass capillaries using doped photocurable ultraviolet resins ^34^. MILL is developed with a standard multiphoton imaging system without the need for specialized 3D laser photolithography platforms ^35^. We further showed that adaptive optics MILL removes spatial optical aberrations in curved thick cylindrical glass capillary tubes. For MILL capillary assays to match with traditional microfluidics, we also describe a rapid hermetical sealing protocol using standard UV adhesive. The UV assisted bonding protocol allows standard silicone fluidic tubes to connect to MILL glass capillary tubes and establish a robust fluidic flow assay. A soft silicone capillary gripper is developed to hold multiple channels for high throughput measurements.

With MILL capillaries, we first recapitulate the dynamics of cell-tissue interactions induced by homogeneous and heterogeneous fluid shear under low FSS (0 – 3.5 dynes/cm^2^) in interstitial tissue and high FSS (0 – 81 dynes/cm^2^) in vasculature. We evaluate these fluid landscapes using cells resident to these tissue (*i.e.* fibroblasts in interstitial tissue and platelets in blood vessels) to map the direction of cell translocating after exposure to heterogenous flow. Fibroblasts adhered by integrin binding and clustering on three-dimensional extracellular matrix (ECM) and initiated cytoskeletal reorganization to alter their physical shape ^36^ and migrate along mechanical and chemical cues. Heterogenous FSS serves as a distinctive mechanical cue (separated from material stiffness), which has previously been observed to stimulate migratory protomyofibroblast phenotype after integrin β1 activation on collagen fibrils ^37–39^. *In vivo* results have shown that fibroblast cells are capable of sensing homogenous shear gradients ^6^.

However, little is known about how fibroblast cells proliferate and differentiate into protomyofibroblasts in irregular interstitial flow conditions. We first established a long-term (12 hrs) live cell imaging protocol using regular capillary tubes that pinpointed the emergence of migratory phenotype resembling protomyofibroblasts ^40,41^ under interstitial fluid flow (< 1 dyn/cm^2^) first observed by Ng *et al* ^39^. Due to lack of molecular markers in protomyofibroblasts (low α-smooth muscle actin (SMA)) ^40^, we focused on migratory responses of fibroblast and alignment of actin cytoskeleton stress fibers ^42^. Using the same interstitial flow rates, we explored heterogeneous fluid flow conditions that mimic tissue microarchitecture or niche environments ^11,43^ and the relationship with fibroblast activation. Using MILL, we further explored how adherent fibroblast cells sense and respond to shear gradients ^44^. Existing fluid shear assays are often conducted on rectangular flow channels that do not faithfully replicate the circular geometry in blood vessels ^45^. We applied MILL to incorporate structures into cylindrical capillary tubes to mimic the appropriate heterogeneous fluid dynamics found in the vasculature ^46^. Using adaptive optics, we showed that a MILL cylindrical tube can take on complex geometries ^47^. To simulate the heterogeneous FSS in cylindrical flow channels, we modelled and designed asymmetrical microstructures ^48^ that mimic small and large stenoses. Since MILL capillary tubes are thin, optically transparent, and smooth, they are highly suited to perform a range of cellular imaging (holographic, confocal, multiphoton and total internal reflectance microscopy) to visualize and quantify several multiple biological markers to assess underlying biological functions.

## Results

### Fibroblast translocation requires integrin engagement and is directly modulated by homogenous FSS

While fibroblasts can proliferate through direct sensing of varying matrix stiffness (1 to 5 kPa ^49^) and ligand concentrations, their ability to respond to FSS within interstitium has so far only been inferred ^37,39,41,42^. To determine the extent to what extent FSS would influence cell translocation, we subjected a murine fibroblast L929 cell line to different levels of laminar fluid shear from 0 to 3.5 dynes/cm^2^ and that mimics fluid shear stress found in interstitial spaces ^39^. We then investigated the influence of ligand binding on cell shape and migratory behavior of fibroblast ^40–42^, where a receptor-ligand adhesion (e.g. integrin-collagen binding) supports increased cell resistance to shear forces over non-specific (e.g. surface charge) interactions. In our experiments, the substrate coating thickness (Poly-L-Lysine (PLL) and collagen fibrils) is a third of the cell thickness, which confers limited force for mechanotransduction. Cells adhering onto PLL or collagen fibrils were exposed to 12 hours of either continuous FSS exposure in transparent capillary tubes or no shear on a coated glass bottom culture dish (see Methods). A portion of cell adhesion could also be accounted by the surface adsorption of fibronectin derived from cell culture serum in both conditions. We quantified migratory patterns and cell shape (determined by cell ellipticity) over 12 hours at 10 second intervals using label-free quantitative phase microscopy (QPM) ^50^ (see Methods), which provides high resolution volumetric morphological profiling. Cell ellipticity (Figure 2 B) i)) quantifies the transition of a cell state from resting to adhesive ^51^. Higher ellipticity values indicate proadhesive cells that are more engaged with ligand-coated surfaces. Under all conditions of shear stress, immobilized collagen appeared to elicit stronger adhesiveness than PLL. The results also suggest that adhesion dynamics can be directly modulated by homogenous FSS alone and is independent of the type of ligand coating. At a shear stress of 1.7 and 2.6 dynes/cm^2^, cell adhesiveness was observed to be low with mean ellipticity of 0.3 and 0.5 for PLL and collagen coatings, respectively (Figure 2 B) ii)). On PLL, cell adhesiveness is highest at 0.9 dynes/cm^2^ with peak mean ellipticity of 0.65. Conversely, cell adhesion on collagen remained high with a peak mean ellipticity of 0.8 at 0.9 dynes/cm^2^. We observed that mean ellipticity reached 0.75 at 3.5 dynes/cm^2^ on collagen. Single cell tracking showed that 3.5 dynes/cm^2^ FSS led to a rapid (<20 sec) expansion of cell perimeter followed by a loss of motion, which we ascribed to cell rupture. Importantly, cell rupture was observed when collagen was the ligand but not PLL (Supplementary Figure S1). In fact, cells adherent on PLL did not rupture over the full 12 hrs. Based on the results in Figure 2 B) ii), integrin adhesion to collagen under FSS ^39^ resulted in a 2- to 3-fold higher surface adhesion when compared with PLL. After cells have adhered to the ligand surface, we tracked the mobility of cells. Under FSS, we repeatedly observed formation of cell aggregates with either ligand coatings. Hence, automated tracking of cell clusters was performed using Trackmate (ImageJ). The results in Figure 2B) iii) shows that cell mobility reached a maximum speed of 9 µm/hr on collagen coating under shear stress of 0.9 dynes/cm^2^ that is around twice the speed when compared with PLL coating. This increase was abrogated at higher FSS (1.7 dynes/cm^2^).

**Figure 2.**
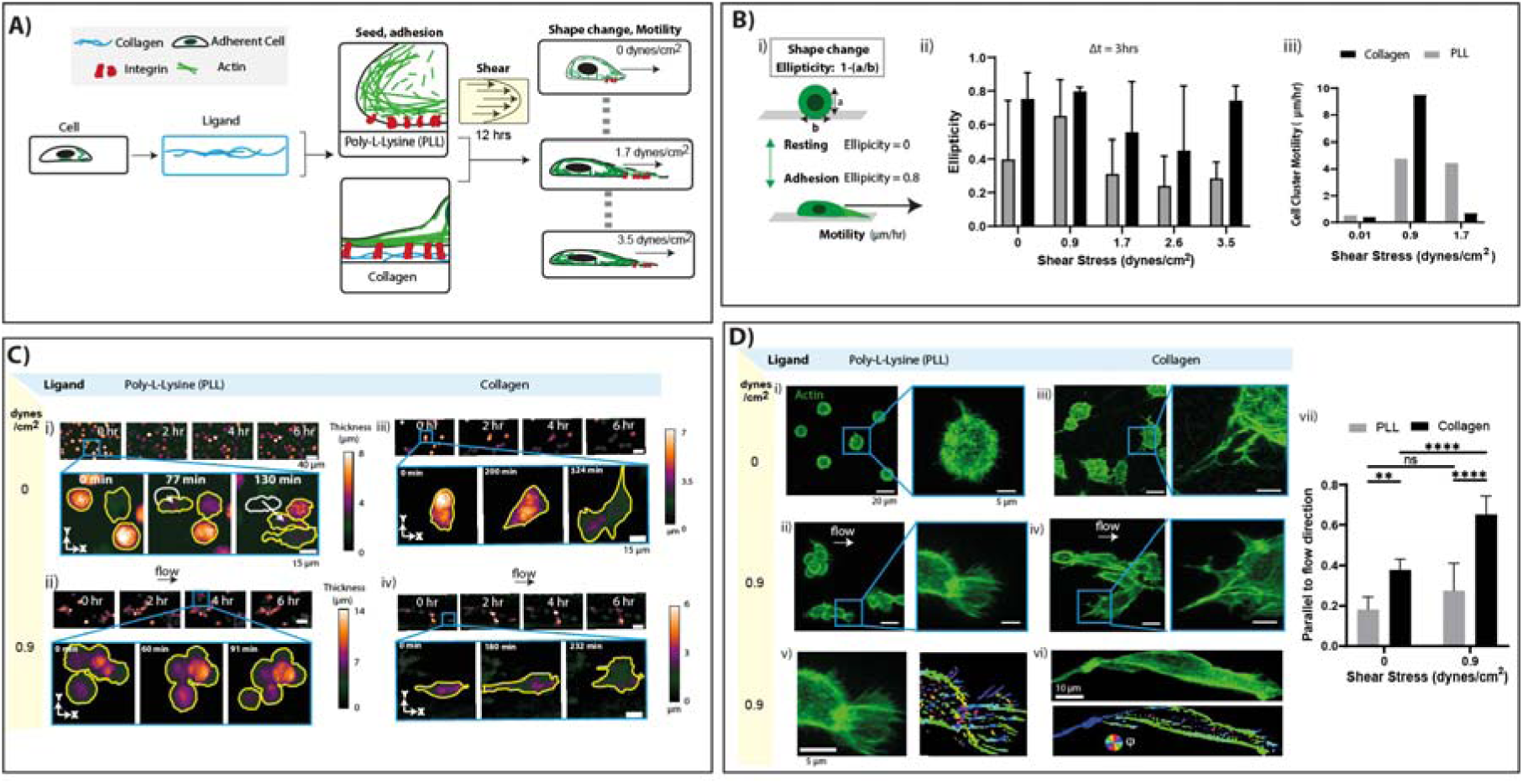
Fibroblast adhesion and proliferation under interstitial, homogenous fluid shear stress: shape change, mobility, and actin reorganization. **A,** Experiment steps to measure cell response (shape change and mobility) along different shear stress and adhesion ligand coatings. Cells (L929, fibroblasts) are first seeded on glass surfaces (dish or capillary tube) that are precoated with collagen or poly-L-Lysine (PLL). A range of homogeneous fluid shear stresses (0 to 3.5 dynes/cm^2^) were imposed over 12 hours period and cell motility monitored by microscopy. **B,** i) Cell adhesion and proliferation are each measured based on cell ellipticity and motility, respectively, using automated cell tracking. ii) Results of cell shape change (ellipticity) subjected to different shear stress (0-3.5 dynes/cm^2^) on collagen or PLL-coated capillaries were tracked over the first 3 hours (n=3 cells). iii) Cell cluster motility at shear stress 00.1, 0.9 and 1.7 dynes/cm^2^. **C)** i) - vi) Time lapse imaging of L929 cells with quantitative phase microscopy over the full 12 hours for selected field of view. Cell cluster motility in B) iii) was measured after cell segmentation (yellow outline). **D)** i) - vi) Confocal maximum intensity projections of actin fiber (F-actin) in L929 cells after 12 hours at shear stress 0 and 0.9 dynes/cm^2^ under PLL and collagen coated glass surface. v) and vi) Segmentation and quantification of actin fiber orientation, alignment along flow direction. vii) alignment of actin parallel to flow (n=4 cells). 0 means completely orthogonal and unity means parallel to flow axis. Scale bars - C) 40 µm and 15 µm, D) 20 µm and 5 µm. Data are the means and SD with D) vi) 2-way ANOVA analysis (**P < 0.01; ****P < 0.001; ns, not significant)

Figure 2C) i) to iv) shows QPM images from a time lapse recording taken at 2-hour intervals. On PLL coating without FSS, most cells (> 80%, n = 37) remained stationary, with a small population that spread and migrated (as shown Figure 2 C) i)), often remaining separated. However, cells exposed to 0.9 dynes/cm^2^ shear stress would spontaneously cluster into multiple discrete aggregates, indicating that intercellular adhesion increased under homogenous FSS. Within each cell aggregate, as shown in Figure 2C) ii), we observed that individual cells would ‘roll’ and organize in the direction of flow. Without flow, cells adhering on collagen-coated capillaries extends membrane protrusions (filopodial) matching half its own cell diameter over several hours (Figure 2 C) iii)). In contrast, as shown in Figure 2 C) iv) and Supplementary Video M1, the first hour of shear exposure (0.9 dynes/cm^2^) to collagen-adherent cells stimulated an increase in motility of up to 9 µm/hr. Together the data indicates shear stress of 0.9 dynes/cm^2^ stimulates maximum adhesion with the highest rate of mobility, a biophysical marker of migratory fibroblast types ^12^. 0.9 dynes/cm^2^ are fluid shear reported in interstitial fluid ^39^ and lymph nodes ^52^, where fibroblasts reside. The presence of collagen and shear are both required to induce a phenotype consistent to proto-myofibroblast formation.

As shown in Figure 2 A), changes to cell shape require actin polymerization to generate tense actin fibers. We quantified actin fiber alignment using F-actin labeling (phalloidin) after exposing cells to fluid shear stress for 12 hrs. We also compared cells that were not exposed to any flow gradient to quantify the influence of fluid shear. On PLL coating, over 80% of the cells (n = 6) retained a spherical shape and actin fibers were randomly arranged (*i.e.,* disordered), shown in Figure 2 D), indicating minimal cytoskeletal rearrangement. Cells adhering to PLL coating that were exposed to homogenous FSS of 0.9 dynes/cm^2^ would generate “fin-like” lamellipodium protrusions whilst remaining weakly aligned, as shown in Figure 2 D) ii). This is consistent with time lapse imaging results from cell shape and motility shown in Figure 2 B) ii) and iii). PLL, in general, shows lower amount of cell adhesiveness and cytoskeleton rearrangements.

On the other hand, cells adhering onto collagen coated surfaces displayed extended protrusions driven by actin polymerization as shown in Figure 2 D) iii) through integrin engagement ^12^. However, under shear stress, membrane protrusions can span over 20 – 30 µm long, as shown Figure 2 D) iv). This result indicates that even at low FSS, actin polymerization can be modulated significantly. Next, we quantified the degree of actin alignment along a chosen axis using morphometric analysis program developed by Lickert *et al* ^51^. In our case, we measure the actin alignment along the flow axis (parallel) as shown in Figure 2 D) v) and vi). A rainbow color code is assigned to the alignment direction. As shown in Figure 2 D) v) and vi), the cells on both PLL and collagen coating were polarized to form actin fibers that align along the flow axis. Actin fibers aligned in parallel to the flow axis will be assigned value close to unity. By examining single cells (n=4) that are not clustered, we observed a two-fold higher alignment of actin to fluid flow on collagen fibers (Figure 2 D) vii), P < 0.001), but no significant increase in in alignment to PLL. Under both absence or presence of fluid shear, we observed ~2-fold increased alignment in cells on collagen compared to PLL coating (P < 0.01). From these results – cell shape, motility, and actin alignment along flow, fibroblast differentiation and translocation are both regulated by integrin engagement and FSS. Additional data on cell viability and time lapse results (Figure 2C) are available in Supplementary Figure S1 and Supplementary Video M1, respectively.

### Homogeneous and heterogeneous fluid shear modulates cell motility and actin bundling differently

In living organs, fluid flow within tissue matrix and microvasculature are mostly heterogenous. Cell-cell interactions are therefore subject to heterogenous shear stresses that can alter signal transductions by a variety of receptor-ligand systems. In contrast to the homogenous FSS, here we asked how heterogenous shear stress modulates fibroblast activation and motility. Based on our findings in Figure 2, we expected heterogeneous shear to form groups of fibroblasts with different levels of adhesion. To investigate this, we need to design and pattern three dimensional physical obstructions within microfluidic channels (Figure 1 B) and 2). To test the throughput of MILL, we generated a 30 µm-thick herringbone structure used to generate turbulent flow for mixing solutions ^53^. The structure consists of 9 herringbones that span the entire width and 1 mm along the length of a square glass capillary (lumen width and height: 200 µm) (Figure 1 B)), where one herringbone took ~9 minutes to complete, demonstrating that MILL achieves millimeter scale structures within 1.5 hours.

Having proven the throughput of MILL, we designed and constructed rectangular obstructions on opposite sides of a capillary (Supplementary Figure S2). The obstructions are three-dimensional microscopic rectangular blocks (W×L: 60 µm × 40 µm) that are spaced at 16 or 24 µm apart with different heights ranging from 6 to 18 µm as shown in Figure 3 A) i). To visualize the lithography structures, we dope the NOA81 optical adhesive with the fluorescent dye rhodamine B. From optimizing dye concentration, we chose a higher dosage of 20 mg/mL, which improved flatness of a lithography layer (Supplementary Figure S3). The flow profile of a standard capillary tube is first calibrated with 1 µm fluorescent microspheres suspended 1× phosphate buffered saline (PBS) and drawn into the MILL capillary tube with an automated syringe pump (Harvard Apparatus). Figure 3 A) ii) shows a cross section of the flow profile indicating a parabolic velocity distribution ranging from 0.2 mm/s to 0.8 mm/s that is typical of laminar flow in a capillary tube. Conversely, in the MILL capillary tubes, we showed heterogenous flow velocity that ranges from a 0.5 to 1.5 mm/s with shear stress ~1.2 dynes/cm^2^ (Figure 3 A) iii)). Fluid flow within the cluster of microscopic obstructions was minimal (<0.5 mm/s) and fastest between obstructions and capillary walls (1.5 mm/s). The same obstructions were also affected by a shear gradient around 100 µm upstream and downstream. The micro-obstruction created a highly heterogenous fluid shear that range from 0.2 to 1.5 dynes/cm^2^, resembling *in vivo* tissue niches within interstitial tissue ^54^.

**Figure 3.**
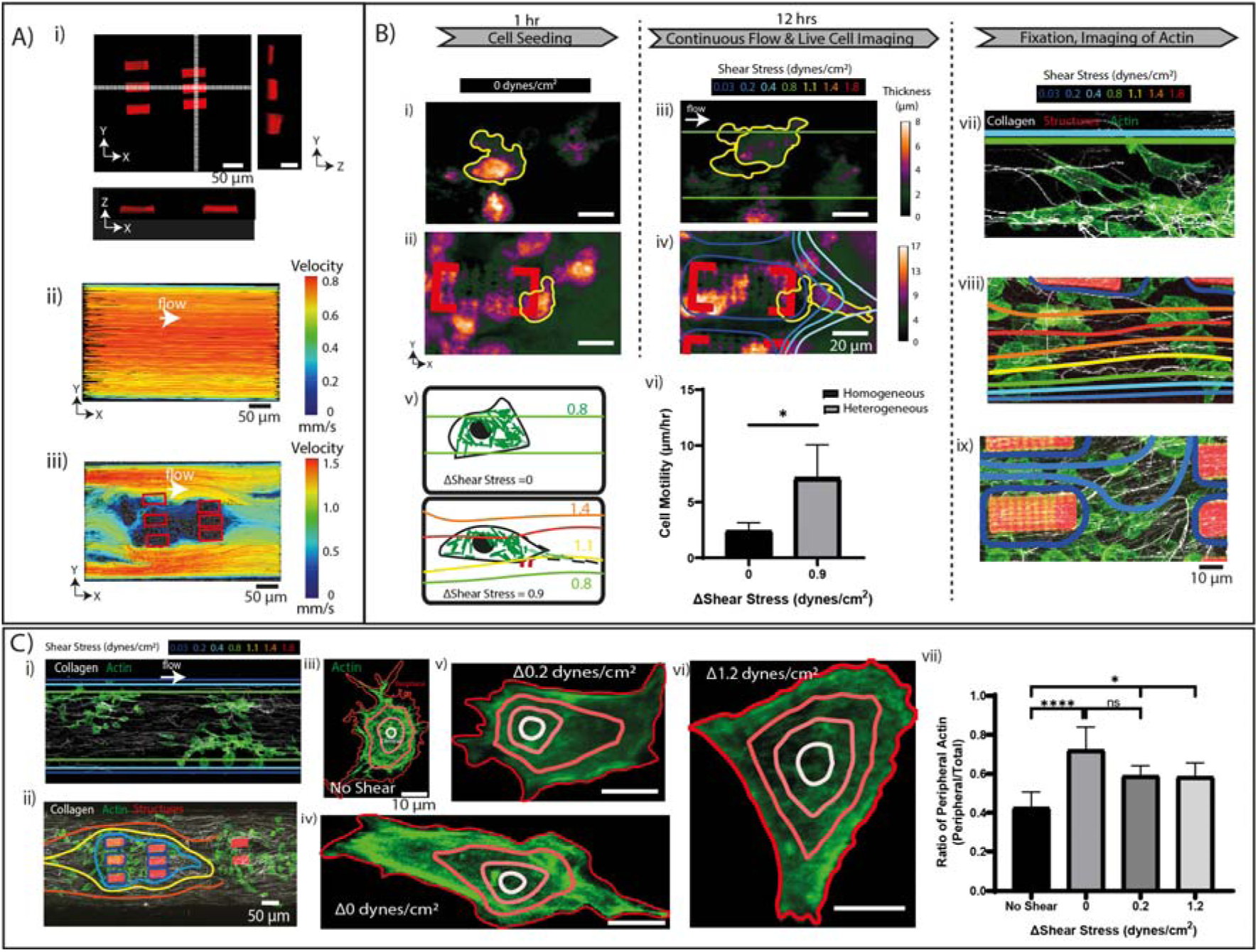
Heterogenous fluid shear stimulates heterogenous cell adhesion, mobility and actin cytoskeleton. **A,** i) orthogonal plots of structures formed by MILL as scanned by confocal microscopy, ii) homogenous and iii) heterogenous flow profile determined by PTV. (**B,** i) and ii)) QPM images of L929 fibroblasts after 1 hour of cell seeding, iii) and iv) after 12 hours of flow to v) determine how heterogeneous shear stress controls cell motility. vi) Quantification of cell motility from time lapse phase imaging (n=3). vii), viii), ix) 2-photon SHG and fluorescence imaging of fixed cells after 12 hours of flow under laminar, homogenous flow rate. **C)** i) and ii) Shows regions with laminar and different shear stress. iii-vi) confocal imaging of actin in L929 cells and segmentation of the cell to vii) identify the proportion of actin at the cell periphery (n=6). Data are the means and SD with B) vi) unpaired t test and C) vii) ordinary one-way ANOVA analysis (*P<0.05; ****P<0.001; ns, not significant).

Next, we determined if the localized regions of shear gradient modulate shape, motility, and actin cytoskeleton of adherent cells. For this, fibroblast cells (L929) were first seeded and left to adhere onto thinly-coated collagen channels for the first hour as shown in Figure 3 B) i) and ii) (collagen coating is shown in Supplementary Figure S2). After which, cells were exposed continuously for up to 12 hours of continuous homogenous and heterogenous fluid shear conditions as shown Figure 3 B) iii) and iv), respectively. Representative images of live cell QPM images shown in Figure 3 B) iii) and iv) identified changes in cell shape and more importantly, mobility. Regions of heterogenous fluid shear showed large changes in cell shape that appear to be sensitive to shear gradient as illustrated in Figure 3 B) v). In Supplementary Video M2, there is a distinct difference in morphology between cells residing within low shear region (~Δ0 dynes/cm^2^) and on margins of high shear stresses at Δ0.9 dynes/cm^2^. We then quantified cell motility under fluid shear change (Δ0 and Δ0.9 dynes/cm^2^) in Figure 3 B) iii) and iv), which showed a significant, 2-fold increase in cell motility in heterogenous fluid shear (Figure 3 B) vi) P < 0.05)). Here, we measured motility in stationary cells that actively formed protrusions (as in B) iv)) and cells that were actively migrating. Next, we used multiphoton imaging to capture the MILL microstructures (Rhodamine B), actin cytoskeleton arrangement (F-actin Phalloidin-Green) as well as collagen fibrils (white) along homogenous and heterogenous shear gradients. We observed that cells organize into distinct clusters around collagen fibers (Figure 3 B) vii-ix)). Importantly, extended actin cytoskeleton appears to coincide with changes in shear stress as shown in Figure 3 B) vii) and viii) and minimal actin reorganization of the cells that resides within rectangular obstruction Figure 3 B) ix) (Supplementary Video M2).

We also correlate the direction in which bundles of collagen fibrils align before and after 12 hours FSS exposure, shown in Figure 3 C) i) and ii). Our results indicate both collagen matrix and adherent fibroblast cells align to fluid shear gradients. Homogeneous FSS modulated fibroblasts appear to exert more force on the collagen matrix than those exposed to heterogeneous FSS Figure 3 C) i) and ii)). This behavior is consistent with the proto-myofibroblast phenotype that has been known to exert forces onto collagen matrix. Because of the high density of clustered cells, it was not possible to quantify single actin fibers as shown in Figure 2 D). Instead, we measured the distribution of fluorescence intensity at different compartments of the cell body (Figure 3 C) iii-vi)). The magnitude of fluorescence intensity indicates bundling of actin fibers in each cell ^55^. Without any FSS stimulus, the actin fluorescence signals appear to be evenly distributed in adherent cells with minimal actin bundling. Adherent cells exposed to either homogenous and heterogenous FSS possessed a significant (P < 0.05) increase in actin bundles along the cell periphery and elongated cell shapes, where homogeneous shear induced 58% higher peripheral actin compared to no shear (P < 0.001). On the other hand, homogeneous FSS (shear gradient of Δ0 dynes/cm^2^, Figure 3 C) vii), stimulated slightly higher increase (~13 %) of actin bundles than heterogenous FSS (shear gradient of Δ0.2 and Δ1.2dynes/cm^2^, Figure 3 C) vii)). These findings indicate that lower heterogenous FSS can lead to an increase in actin bundling and cell adhesion.

### Aberration correction needed for MILL cylindrical capillary tubes

The effects of FSS are most pronounced on cells marginating along the cylindrical walls of blood vessels under flow regimes of high FSS (> 10 dynes/cm^2^). We next determined if MILL could mimic the influence of homogenous and heterogeneous FSS on a cylindrical wall, where an irregular microstructure creates a non-symmetrical shear disruption resembling a vascular stenosis ^48^. We first perform computational fluid dynamic (CFD) modelling to calculate the expected flow rates in a cylindrical tube with inner diameter of 200 µm as shown in Figure 4 A) i) smooth wall and ii) 20% narrowing (stenosis). Along the walls of the 20% narrowing, the fluid flow velocity increased from 0 to 0.3 mm/s within 5 µm from the surface, compared to 24 µm on a smooth wall. Alteration of shear forces along an injured wall not only regulates movement of cells and platelets, it also influences the coagulation pathways involved in thrombus formation ^56^. Cylindrical glass tubes are known to degrade optical performance ^57^. To counter this, we used our Raster Adaptive Optics (RAO) method ^58^ to achieve diffraction limited MILL performance which we termed as Adaptive Optics (AO) MILL. Figure 4 B) i) shows the anticipated written structure of a single pillar without (orange) and with AO (red). We identified degraded optical performance shown in Figure 4 B) ii) and the amount of optical aberrations using RAO as shown in Figure 3 B) iii). Figure 4 B) iv) shows that aberration-corrected laser writing that display sharp edges, measured with QPM. AO MILL enables robust photolithography with a ~15% error in the structure’s dimensions as shown Figure 4 C) v) and vi) in width and thickness. We show that the structure with AO MILL width and thickness was not significantly different from writing on a flat surface with minimal aberration (n=5), demonstrating that AO MILL achieves the writing resolution of the system. We conducted particle tracking velocimetry (PTV), as shown Figure 4 D) at a flow rate of 1 µl/min to experimentally show that the flow velocities match the expected CFD simulation (Figure 4 A)). We also measured the influence of MILL microstructures on the organization of collagen matrix as shown in Figure 4 E) i) and ii). While collagen fibrils adhered to the stenosis structure (red), the overall collagen alignment and distribution across the channel without and with stenosis do not differ significantly (Figure 4 D) iii)).

**Figure 4.**
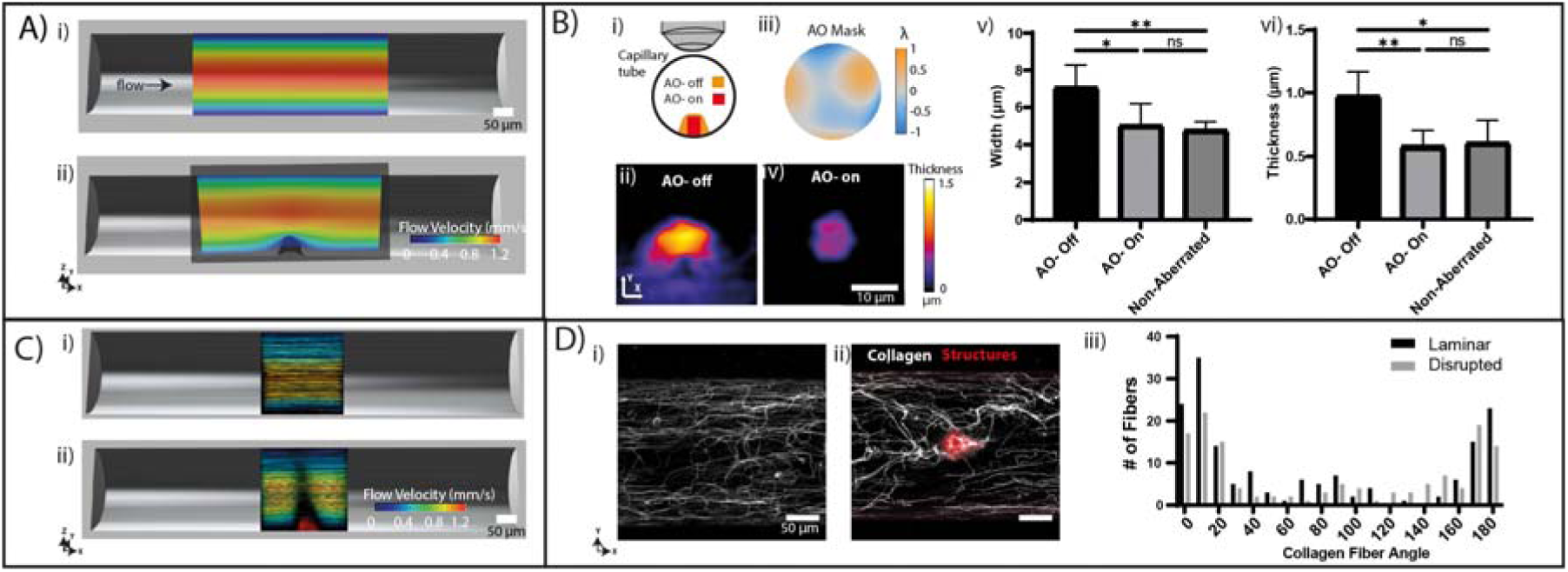
Adaptive Optics Multiphoton Inner Laser Lithography for cylindrical tube. **A,** Non-symmetrical disruption of flow from lithography of asymmetrical structure within a circular capillary with flow velocity determined by CFD simulation. **B**, i) Aberration imposed by cylindrical tube. ii) Lithography structures subjected to aberrations. iii) Identifying aberration mask and iv) lithography structure formed after AO correction and (v) width and (vi) thickness of structures formed with or without AO MILL measured with QPM. Results are the mean and SD of n=5 structures. **C,** i) and ii) Flow velocities across an empty and fabricated AO MILL structure determined experimentally with PTV. **D**, Second harmonic imaging of collagen distribution across a tube under (i) laminar or (ii) heterogeneous shear and (iii) fiber alignment measured after thresholding and segmentation. Data are the means and SD with B) v) and vi) ordinary one-way ANOVA analysis (*P < 0.05; **P < 0.01; ns, not significance).

### Platelet aggregate and move under heterogeneous FSS

The relationship between stenosis and FSS is crucial to our understanding of thrombus formation and blood clotting. Platelets in the vasculature must resist the high FSS force (>10 dynes/cm^2^) by adhering and aggregating to form a thrombus. While platelets are known to be uniquely sensitive to homogenous FSS, the rolling and mobility of adherent platelets around a stenosis is not well studied. Platelet adhesion and shape change remain important biophysical markers for thrombus stability and bleeding disorders ^56^. Thus, we next show that AO-MILL can reveal adhesion and motility of individual platelets as they aggregate onto an adhesive ligand (e.g. collagen). We first coated a thin layer collagen on smooth and AO-MILL cylindrical capillary tubes as shown in Figure 5 A) i) and iv), respectively. Flow velocity of both smooth and AO-MILL tubes are shown in Figure 5 A) ii) and v), where AO-MILL generates heterogeneous shear around the stenosis. Each capillary tube inlet and outlet are connected to a reservoir containing citrated whole blood and a syringe pump, respectively. To form thrombi, the pump draws whole blood across the collagen-coated channel at arterial shear stress of 81 dynes/cm^2^ (shear rate: 1800 s^-1^). To visualize single platelets during thrombi formation, we incubated whole blood with anti-CD42a antibody (ThermoFisher Scientific) conjugated to AlexaFluor 594 to label platelet membrane. To observe thrombus formation in real time and in 3D, we used a video rate multiphoton microscope system ^59^ to record a single volume every second and quantify platelets aggregating. Figure 5 A) iii) and vi) shows a maximum projection of a representative image of several island of thrombus formed after 10 minutes of exposure to fluid shear in two different cylindrical tubes, smooth and stenotic capillary tubes respectively. Results in Figure 5 A) vi) indicates that heterogenous FSS affect the spatial organization of the aggregating platelets. Platelet aggregates appear to organize along the shear direction and within the shear gradient adjacent to the stenosis. The spatial and temporal resolution (1 µm and 20 milliseconds, respectively) ^58^ in our system permits tracking of individual platelets adhering on the coated capillary walls (Figure 5 B)). Tracking results show platelets aggregating ~10 µm away from the stenosis possess an average motility of less than 0.1 µm/s. However, at regions within ~10 µm and downstream to the stenosis, platelets aggregating along the shear stress gradient (55-60 dynes/cm^2^) moved along the flow direction at velocities of up to 2 µm/s (Figure 5 B)). We also captured the 3D trajectory of rolling platelet aggregates as shown in Figure 5 C) i) and ii) and Supplementary Movie M3.

**Figure 5.**
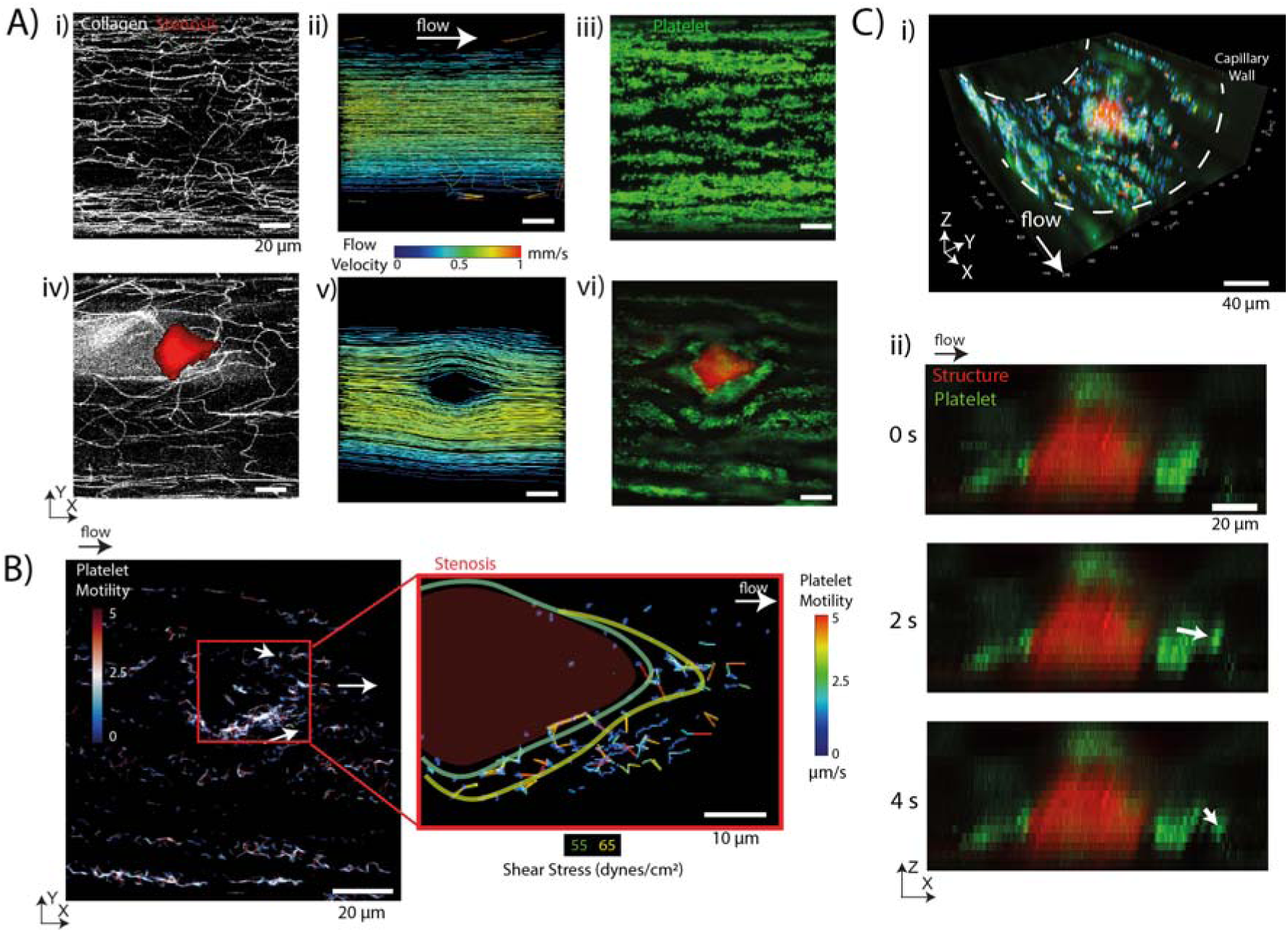
Asymmetrical AO-MILL structure generated heterogenous FSS that increases platelet translocation. **A,** (i) SHG imaging of collagen distribution (white) (ii) flow velocity determined by PTV. (iii) XY slice of platelets adhering on the stenosis, iv) SHG imaging of collagen distribution (white) surrounding an AO-MILL structure (red). (v) flow velocity around AO-MILL structure determined by PTV. **B** tracked to measure platelet motility across the capillary. Arrows indicate direction of platelet translocation. Platelet motility quantified across the capillary in color code and fluid shear stress boundaries are demarcated. **C**, i) Volumetric render of platelet velocities across the stenotic capillary and ii) cross sectional zoomed in image adjacent to the AO MILL structure showing platelet movement (white arrow).

### Degree of laminar flow triggers stenosis triggers platelet adhesion and aggregates

Real time evaluation of platelet adhesion and aggregating profile under different vascular FSS conditions are important functional parameters to determine clotting speed, and platelet dysfunction under a given fluid shear stress. Next, we extended MILL capillary tubes to determine the effects of a larger stenosis with greater heterogeneity in fluid shear stress. We anticipated the FSS gradients from the stenosis and formed platelet aggregate collectively amplify the clotting process to create a large thrombus at the center of the stenosis. To test this, we fabricated a concave structure that mimics an arteriosclerotic vessel ^60^ (Figure 6 A) i)) that disrupts flow velocity by an order of magnitude. Using CFD, we first simulate an input flow of 1 µl/min (1.5 mm/s) that results in a steep gradient of flow velocity of approximately ~ 0.9 mm/s (Figure 6 A) ii)). In contrast with the small 20% stenosis in the previous section, the wall flow velocity with concave structure generates a 3-fold higher acceleration. Using multiphoton microscopy, we measured thrombus formation shown in Figure 6 A) iii) that displays a single large platelet aggregate formed between the concave structure. Using the image, we conducted a second CFD model to estimate the velocimetry along with the thrombus as shown in Figure 6 iv) and a follow up PTV flow measurement in Figure 6) v). The formation of a single large thrombus on an existing stenosis exacerbates the changes in fluid shear to 144 dynes/cm^2^ that would consequentially increase the size of the thrombus and drastically the overall flow gradient. Both simulation and experimental flow velocities confirm a steep acceleration of flow velocity to 1.8 mm/s at the thrombus. For the sake of completeness, we further compared the volume of thrombi formed between homogenous and heterogenous FSS. We observed that homogenous FSS creates a landscape of evenly distributed thrombus across three fields of view spanning across 1 mm in length (Figure 6 B) i)). However, in the case of heterogenous FSS, a single platelet aggregate can reach up to 25 µm thick as shown in the center image of Figure 6 B) ii). The measured distribution of thrombus height also doubled (4 µm) before the stenosis rather than after the stenosis. The overall volume of thrombi in heterogenous flow is around 4-fold higher than homogenous flow as shown in Figure 6 B) iii).

**Figure 6.**
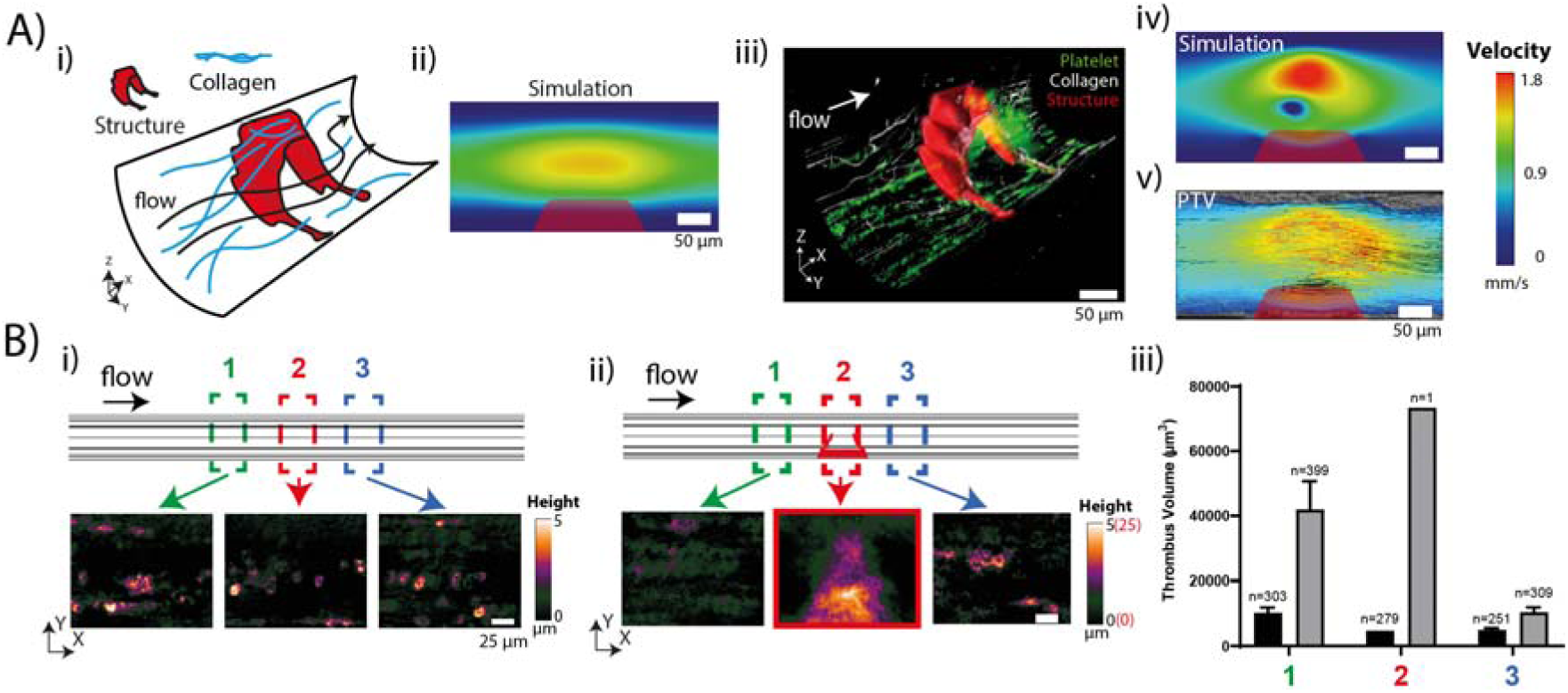
Effect of vascular fluid shear stress increases thrombus formation. **A,** (i) Shear disruption from an irregular AO MILL structure that generates (ii) a heterogeneous flow profile simulated by CFD and promotes (iii) thrombus formation. iv) simulation and (v) PTV after thrombus is formed. **B** QPM imaging of thrombi formed in a capillary **(i)** without and **(ii)** with the AO-MILL structure of formed thrombi upstream and downstream to the stenosis. **iii),** QOM images quantified across three region.

### Increasing throughput of MILL capillaries for thrombosis screening

Microfluidic devices have both throughput and technical advantages over traditional parallel-plate chambers and cone-and-plate viscometer ^61^, such as the precise control of blood flow and the ability to perform multiple experiments with small blood sample volumes. In particular, microfluidics have demonstrated a major role for heterogenous FSS in the initiation and proliferation of platelet aggregation as well as affect antiplatelet therapy on platelet aggregation ^61^. To increase the throughput of these MILL capillaries, we developed a simple PDMS clamp (Figure 7 A)) to mount multiple MILL capillaries and perform homogenous and heterogenous FSS assays in parallel. The role of the PDMS clamp is to secure the capillaries onto a glass slide to prevent sample drifting and rotation, while providing an imaging window and reservoir for immersion objectives. Capillary inlet and outlet tubings are connected and sealed with UV glue. The inner fluid shear profile can be modified with structures using computer aided design and MILL. Following the steps in the previous sections, incorporating multiple capillaries into a single chip allows coating individually or in parallel. Here we focus on collagen coating as shown in Figure 7 B) that is used to evaluate the variation in coating uniformity. Figure 7 C) demonstrates thrombus formed within capillaries generating homogenous and heterogenous FSS using anticoagulated blood treated with an antibody against CD42a, a platelet-specific membrane protein. Using QPM and automated stage scanning, we can rapidly identify and verify the total thrombus volume:area ratio, which is an indication of thrombus height and spread. Figure 7 D) shows the distribution of thrombi formed under shear stress of 81 dynes/cm^2^ for 10 mins in four different capillary tubes, two which have a pair of MILL structures described in Figure 5. These structures were spaced apart by 100 µm, where homogeneous flow is recovered (Figure 4 A) i)).

**Figure 7.**
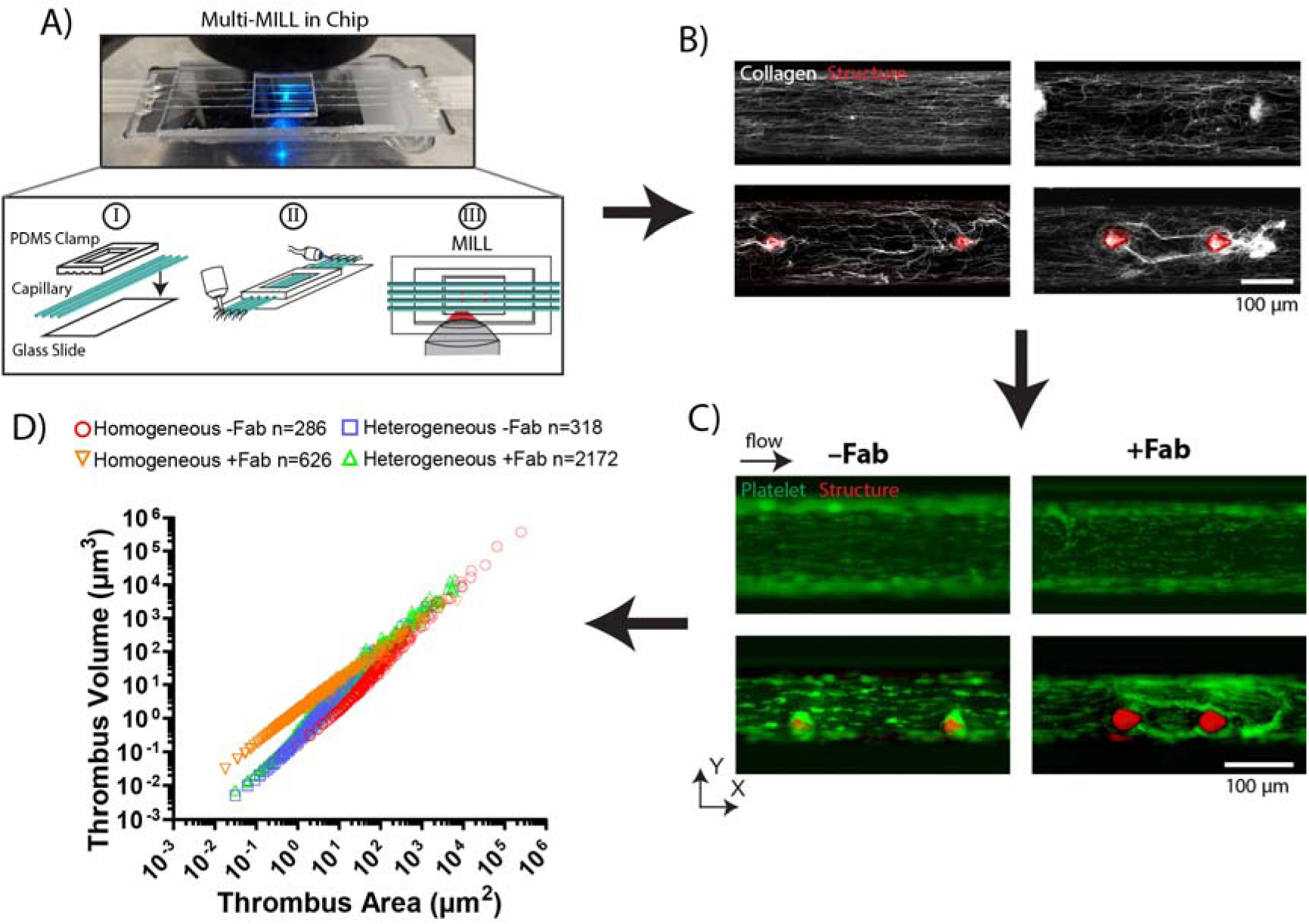
Increasing FSS assay throughput with multiple MILL capillary chip. **A,** Design of a reusable PDMS-based clamp consisting 4 capillary slots and an imaging window, which is adhered onto a glass slide. Inlets and outlets are connected to tubing using an optical UV glue for rapid sealing and structures formed in selected capillaries using MILL. **B**, The collagen coverage is characterized by SHG imaging prior to **C.** thrombus formation under flow in each capillary tube. **D,** Scatter plot of thrombi volume over area as measured by QPM with homogenous and heterogenous flow and with and without GPIV Fab treatment.

We subjected preformed thrombi in two tubes to flow of phosphate-buffered saline alone or containing a monoclonal Fab fragment (10 µg/ml, clone 12A5) to human glycoprotein VI (GPVI) at 81 dynes/cm^2^. GPVI is a platelet-specific membrane protein that binds to collagen and fibrin to mediate platelet adhesion and aggregation^62^. Hence, anti-GPVI Fab treatment will interfere with platelet GPVI-collagen adhesion and remove newly bound platelets at the periphery of the thrombus ^63^. Hence without GPVI Fab treatment, we expect more platelet aggregation and thrombi with larger volume and area than treated thrombi, where platelet aggregates are looser. Our results show that homogeneous flow formed thrombi with area spanning from 10 µm^2^ to 1000 µm^2^ and volumes from 1 µm^3^ to 500 µm^3^. GPVI Fab treatment to preformed thrombi resulted in the majority (76%) of thrombi with area and volume below 10 µm^2^ and 100 µm^3^, respectively. The results also indicated that GPVI-treated thrombi exhibited a higher volume:area ratio, which we predict is due to lower adhesion forces altering thrombus contraction. Under heterogeneous shear, GPVI Fab treatment did not alter thrombi aggregation compared to homogeneous shear, where thrombi volume:area ratio and spreading were similar, with 87% of thrombi below 10 µm^2^. Anti-GPVI Fabs are currently being evaluated as antithrombotic therapeutics as these reagents can interfere with aggregation^64^ or disaggregate formed thrombi ^63,65^ but with minimal bleeding risk ^66,67^. These differences between homogeneous and heterogeneous shear indicate shear disruption could modulate platelet adhesion and hence, alter the efficacy of treatments.

## Discussion

In this study, we demonstrated that MILL outpaces (< 1 hour, Figure 1A) and overcomes the oversimplified laminar flow, 2D geometries of current fabrication techniques. MILL capillaries are accessible *in vitro* systems for studying live cell responses under a uniform (homogeneous) or mixed (heterogeneous) FSS landscape and are tailored for flat or curved flow chambers. These FSS landscapes represent diverse *in vivo* physiological environments from low interstitial FSS in tissue interstitial niche to high fluid shear in vasculature. Importantly, our long-term live cell imaging results showed that heterogeneous FSS landscape produced fibroblasts with increased motility (Figure 3 B) vi)), but reduced cell adhesion (lower peripheral actin) when compared to homogeneous FSS (Figure 3 C)). Heterogenous FSS modulates spatial temporal patterning of platelet aggregates and thrombus formation (Figure 5 and 6). This result reveals fibroblast sensitivity to gradients in FSS. Overall, MILL capillaries can be easily designed to suit conventional CFD modeling and enable quantitative measurement of heterogeneous fluid shear stress on adherent cells. The demonstration of parallel multi-MILL in Chip device also shows that it is possible to tailor different fluid shear stress in a high throughput fashion that could benefit screening of drug efficacy (Figure 7).

### Implication to fibrosis and thrombosis

Our results demonstrate that the FSS landscape regulates cell adhesion in fibrosis and thrombosis. Homogeneous shear increased fibroblast surface adhesion (Figure 2 B) ii)) and actin fiber alignment to the shear direction (Figure 2 D)) by 2-fold in fibroblasts—a response similar to endothelial cells exposed to 15 dynes/cm^2^ of FSS ^68^. However, fibroblasts under heterogeneous FSS resembling interstitial flow (Δ0.2 – 1.2 dynes/cm^2^) were ~ 3 times more motile (Figure 3 B) vi)) and exhibited 50% increase in peripheral actin (Figure 3 C)) compared to homogeneous shear. Likewise, heterogeneous FSS increased platelet translocation (Figure 5 B)) and thrombus contraction by at least 2-folds (Figure 6 B) iii)) compared to homogeneous flow. Therefore, exposing cells to homogeneous or heterogeneous FSS can lead to different conclusions on how fibroblasts and platelets mobilize during fibrosis and thrombosis *in vitro* and *in vivo*. As fluid shear in the interstitium and vasculature are more than an order of magnitude difference, we expect large differences in how cells in both environments sense fluid shear. To gain better insight into how heterogeneous shear controls these cell decisions, future efforts will identify the membrane receptors (e.g. transient receptor potential (TRP) and Piezo protein families) and signaling pathways responsible for sensing FSS in fibroblasts and platelets.

### Implication for organ on chip culture: substrate mechanics and nutrient exchange in glass versus PDMS

An extension of MILL is to generate customized niches for organ on a chip approaches using UV-curable hydrogels ^69,70^ in glass capillaries. *In vivo*, organs consist of stromal tissue with complex a micrometer-scale matrix that produces a heterogeneous FSS landscape. Maintaining the fabrication resolution is important for reproducing this matrix. Hence, we further improve on MILL with adaptive optics to correct for sample aberrations during fabrication. The 2-fold improvement in precision that *AOa MILL* (Figure 4 C)) provides will be important for irregular niches such as bone sinusoids ^11^ and mechanically flexible scaffolds for cardiac cells contraction ^71^. In this study we created a niche mimicking interstitial tissue with heterogeneous FSS using MILL structures spaced apart by 16 or 24 µm (Figure 3 A)). By opting for glass capillaries instead of PDMS, we avoid the non-specific sequestration of hydrophobic molecules in PDMS, which constitute up to 60% of small molecule drugs ^18^, but at the expense of gas permeability ^72^. To support long-term cell culture growth in this niche, we used a HEPES-based buffer that does not require CO_2_ and non-sterile conditions for 12-hour flow experiments (Figure 2 and 3). Restricted gas exchange within the glass capillary tube means fibroblast differentiation and organ development studies lasting >12 hours will require dedicated fluidic setups to maintain nutrient supply, sterility, pH, CO_2_ and O_2_ levels in the culture medium. Existing commercial fluidic pump systems circumvent these demands by integrating a closed fluid circuit within CO_2_ incubators.

### Implication for cell adhesion in metastatic diseases

When fibroblasts were cultured in heterogeneous FSS (*i.e.,* Δ0.2-1.2 dynes/cm^2^, Figure 3 C)), we observed cell translocation against the flow direction (see Supplementary Video M2). This response to flow (rheotaxis) is critical for leukocyte rolling ^73^ in vasculature and metastatic cancer cells ^74^, which can migrate toward a blood vessel and against a flow gradient. Epithelial tumors (e.g. breast cancer) undergo an epithelial to mesenchymal transition (EMT) characterized by gene expression and cell morphology resembling mesenchymal cells ^74^. Cancer-associated fibroblasts (CAF) similarly facilitate cancer migration by remodeling the surrounding ECM ^75^. Elucidating the direct effect of FSS and the contribution of secondary cell mediators to cancer metastasis will be important questions that MILL capillaries can address. A key question is whether cancer cells sense interstitial, heterogeneous FSS to trigger EMT and heterogeneity is regulated by other cells including myofibroblasts ^12^ and macrophages ^76^, which can produce, remodel and pull the ECM.

### Identifying shear conditions that commit cells to divergent cell fates

Adherent and differentiated cells exist in a niche that supports their survival and proliferation, but how does the niche mechanically select for the differentiated cell? MILL capillaries recreate an environment resembling interstitial flow with ‘niches’ of accelerating and decelerating FSS gradients up to Δ1.2 dynes/cm^2^ (Figure 3 A)). *In vivo* shear conditions feature a mix of diverging and converging fluid paths, that is insufficiently described as laminar. Hence, we describe these fluid shear landscapes in MILL capillaries as homogeneous or heterogeneous (Figure 8). We show that these niches regulate fibroblast transition to a proto-myofibroblast state that were up to 3 times more motile (Figure 3 B) vi)) but exhibited 20% less actin bundling (Figure 3 C)) than under homogeneous FSS. Hence, these flow niches act as a mechanical stimulus and generate a shear ‘map’ that regulates fibroblast differentiation. ECM stiffness in cell differentiation is well established ^12^, but evidence shows that FSS is also essential for the differentiation of embryonic stem cells ^13^, endothelial cells ^4^ and fibroblasts ^39^. Future efforts will assess how shear and chemical (e.g. TGF-β) affects the rate of differentiation using molecular markers (e.g. α-smooth muscle actin and cadherin-11) and ECM production (e.g. collagen) as markers of differentiation.

**Figure 8.**
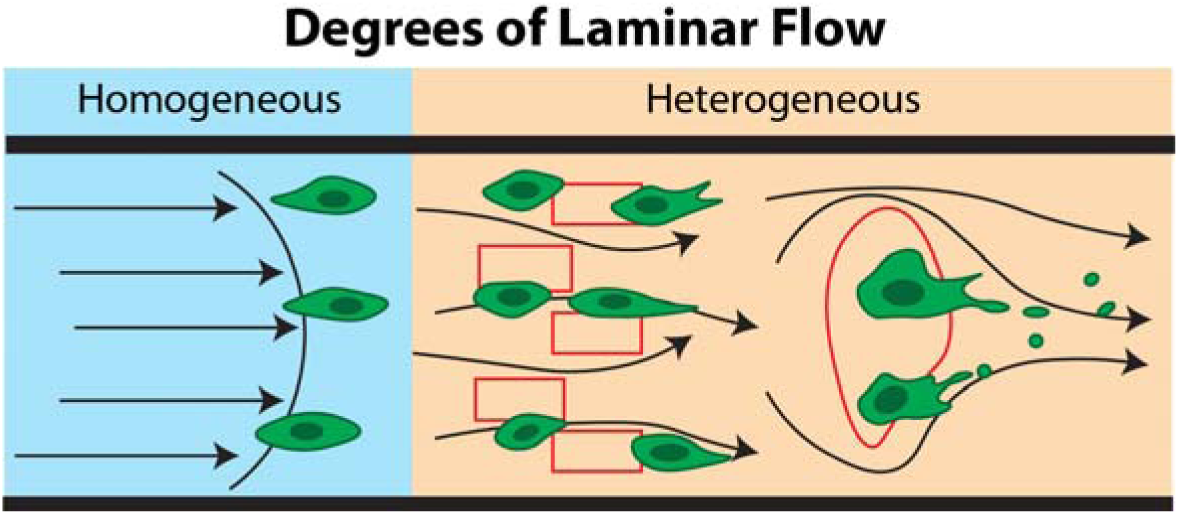
Degrees of laminar flow: homogeneous or heterogeneous flow profiles. Laminar flow is represented in different shear regimes that are classified as homogeneous vs heterogeneous shear distributions, which cells sense and respond to differently.

### Scaling for high throughput screening of pathophysiological responses

Multiwell plates remain the workhorse platform for *in vitro* high throughput screening (HTS) assays of cell behavior. However, current fabrication techniques require hours to replicate the in vivo FSS landscape of stroma and vasculature, that MILL achieves in less than an hour. The fabrication speed sets MILL as a versatile tool to prototype complex *in vivo* FSS landscapes within a single capillary tube, that could not be easily achieved using soft lithography or fused deposition techniques (Figure 2 A). Using MILL we address these restrictions using a standard multiphoton imaging system (Figure 2) to precisely (~15% error) modify FSS within commercial glass capillary tubes (Figure 4 B) v) and vi)). We employed 4 capillary tubes in a single chip assay (Figure 7), which can be scaled for even greater throughput. Incorporating *in vivo* flow parameters to assays will benefit preclinical screening of drug pharmacokinetic and pharmacodynamic properties ^20^. However, as with current HTS methods, machine automation to control flow and reagent input will be required to achieve larger biological scales.

By assembling unstructured and structured capillaries into a fluidic chip (Figure 7), we demonstrate high throughput platelet function testing under laminar (*i.e.,* homogeneous, 81 dynes/cm^2^) and *in vivo* (*i.e.,* heterogeneous, > Δ10 dynes/cm^2^) FSS. Considering heterogeneous FSS abolishes the effect of an antithrombotic drug treatment (Figure 7 D)), coagulation under heterogeneous FSS could be important to infer a patient’s platelet activation status and response to vascular damage or pathology. Current coagulation assays do not account for heterogeneous shear although these represent *in vivo* vasculature (e.g. atherosclerosis ^60^) that are clinically relevant for patients undergoing cardiac surgery. Hence, clotting assays that incorporate heterogeneous flow and shear stress could be used as a screening test for preoperative care and antithrombotic drug selection.

## Conclusion

MILL was shown to recreate both low and high heterogeneous laminar flow rates that are present in porous tissue matrix and blood vessels. We have shown that MILL enables us to study cell adhesion and spreading that is specific to low FSS (0 – 3.5 dynes/cm^2^) landscapes in tissue stroma and high FSS (0 – 80 dynes/cm^2^) landscapes in blood vessels. Furthermore, we demonstrated that MILL fabrication exceeding current 3D fabrication techniques in terms of speed and cost because it can be performed within glass capillaries using routine consumables. This makes MILL well-suited for high throughput cell assays to organ-in-a-chip assays. In our study, we have identified differences of cell translocating under homogeneous and heterogeneous FSS regimes. We first showed that fibroblasts, which reside in porous tissue matrix, respond to low heterogeneous FSS (Δ0.2 dynes/cm^2^), after adhering to collagen fibers, by increasing translocation speed by 3-fold over speed under homogeneous FSS. We also showed that platelets, which initiate repair vessel injury, respond to high heterogeneous FSS (Δ10 dynes/cm^2^) with increased aggregation by 10-fold. We built upon this knowledge of heterogeneous FSS and created multi-MILL in Chip for drug screen that identified a possible effect of heterogeneous shear on antithrombotic drug activity. Taken together, the use of MILL would expand the exploration of heterogeneous physiological shear environments for other biological cells type i.e. macrophages ^8^, cancer cells ^6,77^. MILL assays can therefore capture *in vivo* cell responses under both physiological and pathological fluid shear landscapes and so can be used as pharmacological assays without resorting to complex microfluidic systems ^78^.

## Methods

### Preparation of Capillary Chips

For all experiments, borosilicate capillaries were cut to a length of 35 mm (rectangular tubes) or 85 mm (circular tubes) and mounted onto 75 mm × 26 mm microscopy glass slides. Rectangular capillaries (W × H; 0.3 mm × 0.1 mm, VitroCom, Mountain Lakes, NJ, USA) were secured onto the glass slide using NOA81 UV adhesive (Norland Products, Cranbury, NJ, USA) with a UV curing system (365 nm, Thorlabs, Newton, NJ, USA) at 20% intensity for 5 seconds. To secure circular tubes (ID: 0.2 mm, VitroCom) onto glass slides, a mold formed with circular capillaries was used to create a PDMS clamp as shown in Figure 6 A). Capillary ends were inserted into Tygon inlet and outlet tubing (ID: 0.51 mm, Saint-Gobain, Courbevoie, France), which was sealed with NOA63 UV adhesive (Norland Products) and UV-cured at 20% intensity for 30 seconds.

### MILL and RAO MILL

UV adhesive for MILL was prepared by dissolving Rhodamine B (20 mg/mL, Townson & Mercer, Australia) in NOA81 UV adhesive (Norland Products) by manual mixing and through a suspension mixer for 2 hours and then injected into borosilicate capillaries with a 1 mL syringe. MILL was performed using a custom-built polygon scanning microscope ^59^ with a Ti-Sapphire pulse laser (Spectra Physics, MKS Instruments, Inc., Andover, MA, USA) tuned to 810 nm with a pulse width of 100 fs and repetition rate of 82 MHz. We used an average laser power of 22 mW and 1.2 mW after the 20× water-immersion objective lens (W Plan Apochromatic, 1.00 NA, Zeiss, Germany) for MILL and imaging, respectively. Patterning of microstructures using 2PP was achieved by restricting the scanning range to 6 µm × 4 µm at the image plane and translating the scanned region laterally with galvo mirrors to form the structure. Depth of the MILL structures was controlled with the sample stage (3DMS, Sutter Instrument, Novato, CA, USA) and tested for lithography precision (see Figure 3 B) v-vi)). MILL was also replicated on an Olympus FVMPE-RS multiphoton microscope controlled by the Fluoview software (data not shown). Unpolymerized adhesive was washed out with acetone and ultrapure water (Milli-Q, Merck, Rahway, NJ, USA).

RAO was performed as previously described ^58^. Briefly, the sample aberrations were determined at a depth matching the height of the structure to be written. The deformable mirror is stepped through the first 11 Zernike modes (excluding tilt, piston and defocus) and amplitudes using the fluorescence signal as feedback. The identified Zernike mask was then applied to the entire MILL structure.

### Expression and Purification of Anti-GPVI Fab

Mouse anti-human GPVI monoclonal antibodies were generated (The WEHI Antibody Facility, Melbourne, Australia) and isolated from hybridoma supernatant as previously described ^64^. Antibody in the hybridoma supernatant was purified by passing through a column of DEAE Affigel blue (Bio-Rad, Sydney, NSW, Australia), dialysed into a Tris-saline buffer (Tris 10 mM, NaCl 150 mM, pH 7.4) then affinity purified on a protein A sepharose column (GE Healthcare, Chicago, IL, USA). Bound antibody was eluted with 0.1 M Glycine pH 2.4 and neutralised with 1 M Tris pH 8.5. Purified antibody was then dialysed into phosphate buffered saline (NaCl 137 mM, KCl 2.7 mM, KH2PO4 1.8 mM, Na2HPO4 10 mM, pH 7.4). Fab fragments were generated from these antibodies using a Fab preparation kit (Pierce Biotechnology, Rockford, IL, USA) according to manufacturer’s instructions.

### Cell Culture and Shear Assays

All cell culture reagents were purchased from Thermo Fisher Scientific (Waltham, MA, USA). Murine L929 fibroblasts (ATCC, Manassas, VA, USA) were maintained in low glucose (1 g/L) DMEM supplemented with 10% fetal bovine serum, L-glutamine (4 mM) and pyruvate (1 mM) at 37°C and 5% CO_2_. Cells were split 1:6 at 80% confluence.

For shear assays, fibroblasts in growth phase were detached with trypsin-EDTA (0.05%) and centrifuged at 300 × *g* for 5 minutes. Supernatant was removed and cells were resuspended at 2.5 ×10^5^ cells/mL in 1 mL of HEPES-Krebs buffer (120 mM NaCl, 22 mM HEPES, 4.6 mM KCl, 1 mM MgSO_4_, 155 µM Na_2_HPO_4_, 412 µM KH_2_PO_4_, 5 mM NaHCO_3_, 1 g/L glucose, 1.5 mM CaCl_2_, pH 7.4) supplemented with 10% (w/v) fetal bovine serum. Cells were seeded on glass-bottom dishes (no FSS condition) or flowed into the capillary at 2.5 mL/min (with FSS) on a stage heated to 37°C and allowed to settle for 1 hour. Krebs-HEPES buffer was then flowed through at different velocities equating to shear stress values of (0 – 3.5 dynes/cm^2^) for up to 12 hours and numbers and behavior of adherent cells were monitored with label-free quantitative phase imaging.

### Capillary Thrombus Assay

Circular capillaries (ID: 0.2 mm, 65 µm wall thickness) were precoated with Type-I Collagen (HORM, Takeda Austria GmbH, Linz, Austria) for 1 hour and washed with 1× PBS at a flow rate of 80 µl/min for 5 minutes. Whole blood was collected in citrated (3.2% w/v, Sigma-Aldrich, St. Louis, MI, USA) saline and incubated for 30 min with anti-CD42a antibody (1/100 dilution, clone FMC-25, Thermo Fisher Scientific) conjugated to AlexaFluor 594. The blood was flowed using a syringe pump (PhD Ultra, Harvard Apparatus, Holliston, MA, USA) on withdraw mode and at a shear stress of 1800 s^-1^ for 10 min, followed by PBS or anti-GPVI Fab (10 µg/ml in PBS, clone 12A5) for another 10 min. Thrombi were fixed with 4% paraformaldehyde (Sigma-Aldrich) for 10 minutes and then washed with PBS for 10 minutes under 80 µl/min flow. All blood samples were taken with consent from each donor and approved by the ANU research ethics office (2022/372).

### Multiphoton Imaging and Deconvolution

Rapid live imaging of thrombi formation was performed using a custom-built polygon scanning microscope ^59^ at 20 frames per second and rapid volumetric scanning (21 Z-slices per stack) was achieved by tuning the defocus Zernike amplitude on a deformable mirror conjugated to the back focal plane of the objective. Fluorescence emission was measured through a detection unit consisting of 2 dichroic mirrors (FF-562-Di02, Semrock, IDEX Health & Science, Rochester, NY, USA), emission filters (FF01-514/44 and FF01-624/40, Semrock) and three GaAsP photomultiplier tubes (H7422-40, Hamamatsu, Hamamatsu City, Japan).

SHG and actin imaging were performed with an Olympus FVMPE-RS multiphoton microscope and a 25× water-immersion objective lens (XL PLAN W MP, 1.05 NA, Olympus, Tokyo, Japan). The excitation laser was tuned to 900 nm at a laser power setting of 15% and emission signal detected through an FV30-FVG filter set (dichroic: SDM475, emission filters: BA410-455 and BA495-540) and 2 GaAsP photomultiplier detectors. Volumetric images were obtained with a resonant mirror scanning at 15 frames per second with 8-line averaging.

A customized ImageJ macro based on the CLIJ2 plugin ^79^ was written for deconvolving volumetric images obtained from the polygon 2P microscope. Images were deconvolved using the Lucy-Richardson algorithm with 20 iterations and an experimental PSF obtained from imaging of 1 µm yellow-green, fluorescent beads (Polysciences, Warrington, PA, USA).

### Confocal Imaging

High resolution confocal imaging of actin was performing with a Leica SP5 microscope with 488 nm and 561 nm excitation lasers and 63× oil immersion objective (HCX PL APO, 1.4 NA, Leica Microsystems, Wetzlar, Germany). Fluorescence emission was measured using hybrid GaAsP detectors and emission cutoffs of 502-545 nm for Actin Green and 584-660 nm for rhodamine B. Scanning was performed with a 4-line averaging and pinhole set to 1 Airy Unit.

### Label-free Quantitative Phase Microscopy

Quantitative phase microscopy was performed using a custom-built imaging system with a 514 nm laser (OBIS 514nm LS 20mW Laser, Coherent Inc., Santa Clara, CA, USA). The laser coupled to an optical fiber, which was then split into an object and reference path. Light from the object path is collimated and passed through the sample and imaged through a 20× microscope objective (UCPlanFL N, 0.4 NA, Olympus). The transmitted light is interfered with the reference light and was captured by a CMOS camera (BFS-U3-32S4, Teledyne FLIR LLC, Wilsonville, OR, USA). The phase information was reconstructed by an open-source MATLAB program (DHM_MATLAB_ANUAOLAB, V4.0. Source code available at https://github.com/PurelyWhite/DHM_MATLAB_ANUAOLAB) and visualized in Fiji (ImageJ).

### Computational fluid dynamics simulations

Flow simulation was performed using COMSOL Multiphysics software (Burlington, MA, USA). Briefly, blood or plasma were represented as a Newtonian fluid with the viscosity of water (1 mP s). The mass inflow of 1 mg/min (corresponding to 1 µl/min) in a cylindrical vessel of 200 µm diameter and 800 µm length was considered as boundary condition for the inlet and zero pressure for the outlet. No-slip boundary conditions were chosen for the rest of the system boundaries and laminar flow module with incompressible fluid approximation was used for numerical analysis of a stationary Navier-Stokes equations solution. Magnitudes of flow velocities were extracted in a regular grid with 1 µm spatial resolution. 3D reconstruction of the stenosis geometry was performed with Fiji (ImageJ) capability to render a wavefront object from a volumetric scan of a MILL structure. The triangulation parameters were kept default (threshold 50, resampling factor 2). The Wavefront objects were imported into the Autodesk Fusion 360 (Autodesk, San Rafael, CA, USA) as the mesh object. Each mesh object was cleaned from noise and smoothed. An array of surfaces was created to obtain a set of projection sketches as an intersection of a mesh body with a surface. The capillary object was created as a transitional shape between projection border profiles and saved in the appropriate format for the COMSOL Multiphysics import.

### Particle Tracking Velocimetry

1 µm yellow-green beads (Polysciences) were flowed through a capillary tube by pulling at 250 nL/min (rectangular capillary) or 1 µl/min (circular capillary) and imaging was performed by resonant scanning in the Olympus multiphoton (scanning at 22 Hz) or polygon scanning microscope (25 Hz) with AO, respectively. Beads were tracked using TrackMate ^80^ to obtain the velocity and trajectory using the Kalman filter for homogeneous flow or simple LAP tracker for heterogeneous flow, and a maximum spot displacement threshold of 20 µm.

### Measurement of Collagen and Actin Morphometric Phenotyping

Collagen fiber alignment was measured using the MATLAB code CT-FIRE developed by Liu and colleagues ^81^. For this, a volumetric SHG scan was analyzed and collagen angle at each Z-slice quantitated.

Actin alignment of high-resolution confocal images was measured using a MATLAB (MathWorks, Natick, MA, USA) script written by Lickert *et al.* for cell segmentation and actin fiber measurements ^51^. Quantitation of peripheral actin density was performed using the Measure Object Intensity Distribution module in Cell Profiler ^82^.

### Cell and Platelet Tracking

Time lapse videos of L929 cells and platelets were tracked using TrackMate. Individual platelets were identified using the Laplacian or Gaussian detector. Due to variations in the cell morphology, cell segmentation was performed using the Thresholding detector or CLIJ2 Voronoi Otsu Labeling, dependent on which distinguished individual or clustered cells more accurately. To remove misidentified tracks, each result was optimized for maximum track length, and mean directional change. The selection of filters on spots and tracks with the relevant parameters setting were tuned for each dataset to achieve the best tracking performance. Analysis of cell motility and morphology was done based on the tracking data provided by TrackMate.

### Measurement of Cell Ellipticity

Reconstructed quantitative phase images with height information of the cells were plotted with the Fiji function ‘3D Surface Plot’ with a custom-developed user interface for parameter control (viewing angle in XY plane, viewing angle in XZ plane, perspective, smoothing and global height modification). The self-developed user interface was used to provide the side projections (XZ projection) of the cells and to generate a mask of the cell surface. The surface was fitted to an ellipse, where the height and length of the cells were used to calculate cell ellipticity according to the definition of first flattening (f = 1-b/a), where ‘b’ is the longer axis and ‘a’ is the shorter axis.

### Statistical Analysis

Data were analyzed using Prism (version 9.3.1, Graphpad Software, San Diego, CA, USA). Ordinary one-way and two-way ANOVAs were performed with Tukey’s test. Unpaired parametric t-tests were performed with P-values calculated using two-tailed analysis.

## Author Contributions

W.M.L. initiated, developed and supervised the project. Y.J.L. led the project, performed experimental work and carried out the analysis of the results. J.Z. prepared and performed cell experimental work. H.L. and T.X. assisted in lithography. H.L. assisted in image analysis and deconvolution. Y. L. built the AO system. Z. Z. advised on software development of phase imaging. S.M.H. and E.E.G. advised on platelet experiments and S.M.H. prepared GPVI Fabs. I.C. and D.N. carried out flow simulation. W.M.L. and Y.J.L. wrote the manuscript with input from all authors.

