## Supplementary Information for "Patterning High to Low Heterogenous Fluid Shear Stress Landscapes with Multiphoton Inner Laser Lithography (MILL) for Live Cell Adhesion and Translocation"

### Supplementary Videos

**Supplementary Video M1. 12-hour live imaging of L929 fibroblasts on PLL or collagen with or without homogeneous shear.** Fibroblasts were seeded on glass-bottom cover dishes or rectangular capillary tubes coated with PLL or collagen, then exposed to 0 or 0.9 dynes/cm$^2$ shear for 12 hours. Changes to cell thickness were measured with quantitative phase microscopy. Translocating (*) cells are indicated in the video.

**Supplementary Video M2. L929 fibroblast response to homogeneous or heterogeneous shear.** Fibroblasts in collagen-coated rectangular capillary tubes were subjected to homogeneous (no structures) or heterogeneous (with MILL structures) shear stress for 12 hours and imaged with quantitative phase microscopy. Lines indicate regions of different fluid shear stress. Translocating (*), resting (^) and extending/retracting (+) cells are indicated in the video.

**Supplementary Video M3. Real-time volumetric imaging of thrombus formation under homogeneous or heterogeneous shear.** Whole blood was flowed through a collagen-coated circular capillary tube with or without MILL structure for 5 minutes at a shear stress of 81 dynes/cm$^2$. Thrombus formation was imaged in real-time (1 volume/second) by multiphoton microscopy.

**Supplementary Video M4. Real-time volumetric imaging of thrombus formation under large fluid shear disruption.** Whole blood was flowed through a collagen-coated circular capillary tube with large stenotic MILL structure for 10 minutes at a shear stress of 81 dynes/cm$^2$ and thrombus formation was imaged in real-time (1 volume/second) by multiphoton microscopy. Arrow indicates the direction of a translocating thrombus (*).

### Supplementary Figures

**Figure S1- Shear-induced changes to L929 morphology and viability. A,** L929 cells were cultured (i) on a glass bottom dish or (ii) under FSS within a glass capillary. (**B,** i) DHM imaging of cells on surfaces with/without PLL or collagen coating and under increasing FSS. (ii-iii) Cells were segmented and the number of cells on (ii) PLL- or (iii) collagen-coated surface was measured over 3 hours.

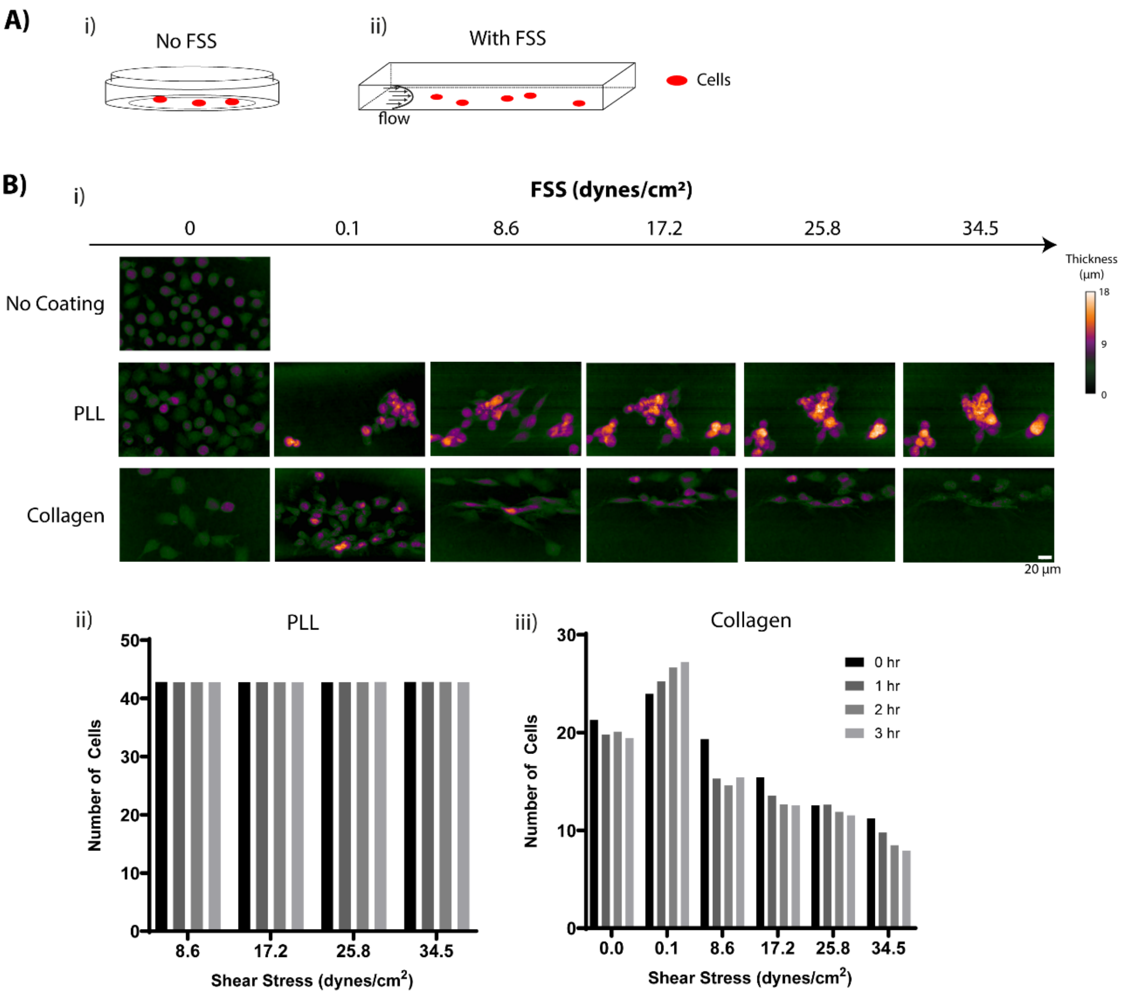

**Figure S2. Measurement of collagen coating thickness.** SHG imaging of capillaries **(A)** without or **(B)** with MILL structures. (i) XY maximum projections from volumetric SHG imaging after coating of capillaries with collagen. (A, ii and B, ii-iii) shows a zoomed in field of view boxed in (i) and (A, iii and B, iv-v) YZ projections across the indicated region (dashed line). **C,** Bar plot of collagen coating thickness measured from the YZ projections (dashed yellow line).

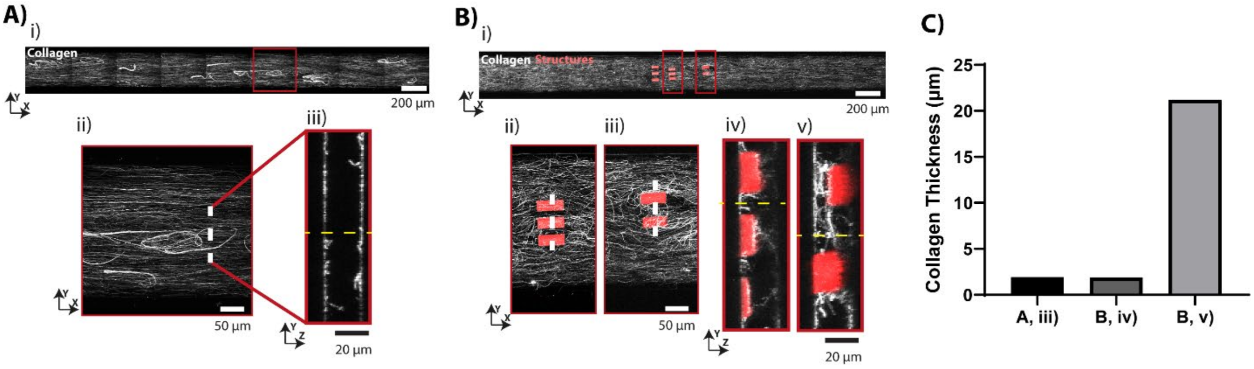

**Figure S3**. **Rhodamine B facilitates MILL structure prototyping. A,** Lithography of a 10 µm-thick window pattern on a glass coverslip using NOA81 glue doped with different concentrations of rhodamine B and imaged with DHM. (ii) Plot of structure height across the line profile taken from i, dashed line. **B,** Confocal imaging of window structures doped with 10 and 20 mg/mL of rhodamine B.

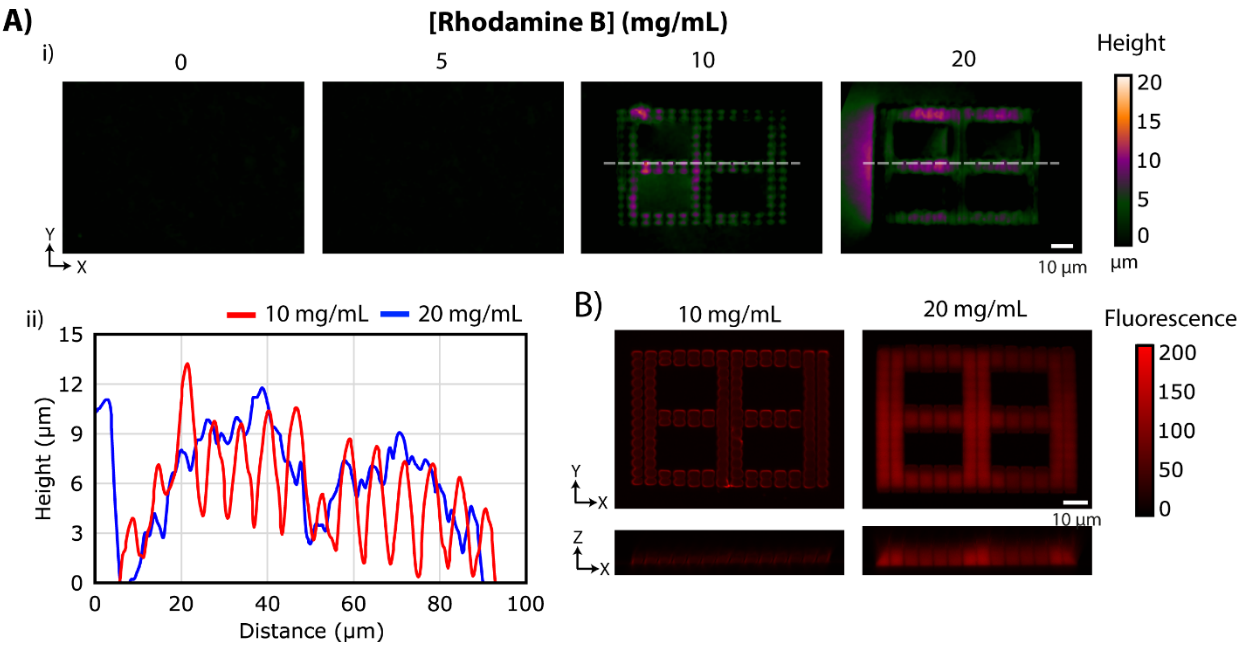
